# Sensory- and memory-related drivers for altered ventriloquism effects and aftereffects in older adults

**DOI:** 10.1101/2020.02.12.945949

**Authors:** Hame Park, Julia Nannt, Christoph Kayser

## Abstract

The manner in which humans exploit multisensory information for subsequent decisions changes with age. Multiple causes for such age-effects are being discussed, including a reduced precision in peripheral sensory representations, changes in cognitive inference about causal relations between sensory cues, and a decline in memory contributing to altered sequential patterns of multisensory behaviour. To dissociate these putative contributions, we investigated how healthy young and older adults integrate audio-visual spatial information within trials (the ventriloquism effect) and between trials (the ventriloquism aftereffect). With both a model-free and (Bayesian) model-based analyses we found that both biases differed between groups. Our results attribute the age-change in the ventriloquism bias to a decline in spatial hearing rather than a change in cognitive processes. This decline in peripheral function, combined with a more prominent influence from preceding responses rather than preceding stimuli in the elderly, can also explain the observed age-effect in the ventriloquism aftereffect. Our results suggest a transition from a sensory- to a behavior-driven influence of past multisensory experience on perceptual decisions with age, due to reduced sensory precision and change in memory capacity.

## 1. INTRODUCTION

One of the commonly known features of healthy aging is losing the ‘sharpness’ – such as the precision of sensory perception (Dobreva et al., 2011; Lindenberger & Baltes, 1994; Otte et al., 2013; Salthouse, 1996), that of memory (Salthouse, 2010, 2019), or the swiftness of responses (Falkenstein et al., 2006; Jones et al., 2019). Such changes in perception are accompanied by alterations in brain structure or brain activity with age (Henry et al., 2017; Henschke et al., 2018; McNair et al., 2019), suggesting that aging may affect both early sensory and higher level cognitive processes (Diaconescu et al., 2013; Dully et al., 2018; Henry et al., 2017; McNair et al., 2019; Nunez, 2015; Zanto & Gazzaley, 2014). Unravelling the mechanisms by which perception changes with age becomes particularly challenging when behavior relies on the combination of multiple sensory or behavioral attributes, such as during multisensory integration or the adaptive trial-by-trial recalibration of perception based on previous multisensory experience (also known as after-effects). Indeed, age-related changes in multisensory behavior could result from a number of sources, including peripheral sensory processes, decision strategies or reduced resources such as attention or memory. Hence, pinpointing whether and why multisensory perception changes with age has remained difficult (de Dieuleveult et al., 2017; Dully et al., 2018; Freiherr et al., 2013).

Previous studies have described a wide range of apparent changes in multisensory behaviour with age (de Dieuleveult et al., 2017; Freiherr et al., 2013): older adults seem to benefit more from multisensory information during speeded detections (Diaconescu et al., 2013), they are influenced more by task-irrelevant distractors (Dobreva et al., 2012), have wider multisensory temporal binding windows (Bedard & Barnett-Cowan, 2016; Chan et al., 2014b; Colonius & Diederich, 2011; Diederich et al., 2008), and sacrifice response speed to preserve accuracy (Jones et al., 2019). Furthermore, the adaptation to audio-visual synchrony and the recalibration during temporal binding are both attenuated in older adults, possibly as a result of memory decline (Bedard & Barnett-Cowan, 2016; Chan et al., 2014a; Dobreva et al., 2012; Noel et al., 2016; Vercillo et al., 2017).

Despite these widespread changes in multisensory perception with age, the underlying mechanisms remain debated and multiple causes have been suggested. First, changes in peripheral sensory processes (e.g. hearing loss) can reduce the reliability of the sensory evidence provided by a one modality. This effectively reduces the influence of this modality on behaviour while increasing that of another during multisensory integration (Dobreva et al., 2012; Lindenberger & Baltes, 1994; Trelle et al., 2019; Tye-Murray et al., 2016). Such a peripheral origin predicts that changes in multisensory behavior are accompanied by detectable changes in unisensory performance. Second, changes in cognitive strategies can affect how causal relations are inferred from disparate evidence and thereby shape whether and how these are fused for a perceptual judgment (Baum & Stevenson, 2017; Diederich et al., 2008; Laurienti et al., 2006; Mozolic et al., 2012). These processes of sensory causal inference have been linked to parieto-frontal brain function and can be captured using Bayesian models of causal inference (Cao et al., 2019; Körding et al., 2007; Rohe & Noppeney, 2015a; Wozny et al., 2010). Such an origin should hence be reflected by corresponding Bayesian models applied to behavioural data. And third, changes in working memory can influence the degree to which previous sensory information shapes behaviour in consecutive trials (Allred et al., 2016; Crawford et al., 2016; Dobreva et al., 2012). Brain structures involved in working memory have been recently implied in multisensory trial-by-trial recalibration (Park & Kayser, 2019) and thus memory-related contributions to changes in multisensory behaviour with age should reflect in recalibration paradigms.

The focus of this study was to test and understand age-related changes in a well-established model paradigm for multisensory perception: the spatial ventriloquism task. We compared behavioural performance in two frequently studied multisensory response biases between young (18 ~ 35 years) and older (62 ~ 82 years) healthy participants obtained in the same paradigm: the within-trial integration of multisensory information (the ventriloquism effect) and the rapid trial-by-trial recalibration of unisensory perception based on a previous multisensory stimulus (the so-called immediate ventriloquism aftereffect)(Wozny & Shams, 2011b). These two multisensory biases capture distinct aspects of how multiple stimuli can interact: the ventriloquism effect allows characterizing causal inference processes about discrepant multisensory information within a trial as well as the relative weighting of unisensory cues. On the other hand, the ventriloquism aftereffect characterises the persistence of multisensory information over time and its influence on subsequent decisions. We used both model-free and model-based analyses to isolate and quantify components shaping these biases to better understand the age-related changes in these two ventriloquism biases.

## 2. METHODS

### 2.1. Participants and screening procedures

24 healthy right-handed younger adults (YA, 9 males, mean age 23.5 years, range 18 ~ 35 years) and 24 healthy right-handed older adults (OA, 7 males, mean age 69.0 years, range 62 ~ 82 years) participated in this study. The sample size was determined based on previous studies using similar experimental protocols (Jones et al., 2019) and recommendations for sample sizes in empirical psychology (Simmons et al., 2011). All participants submitted informed written consent. The YA had self-reported normal vision and hearing and indicated no history of neurological diseases. OA’s were screened for normal vision and hearing: pure-tone audiometric thresholds were obtained at 500 Hz, 1000 Hz, and 2000 Hz and individuals with average thresholds higher than 30 dB for either ear were excluded (mean ± SD thresholds for included participants: 14.1 dB ± 6.4 dB, 13.8 dB ± 6.8 dB for the left and right ear respectively). The visual acuity was 20/25 or 20/20 for all OA participants. OA were also tested on the Montreal Cognitive Assessment (MoCA) (Nasreddine et al., 2005) and all OA’s scored above 26, indicating no cognitive impairment (mean ± SD: 28.8 ± 1.56). Data from two YA (both females) had to be excluded as they were not able to perform the task correctly. One OA did not pass the hearing test and two did not pass the spatial hearing test (see below; both females). Therefore, data are reported for a sample of 22 YA and 21 OA.

### 2.2. Stimuli

The acoustic stimulus was a 1300 Hz sine wave tone (50 ms duration) sampled at 48 kHz and presented at 64 dB r.m.s. through one of 5 speakers (Monacor MKS-26/SW, MONACOR International GmbH & Co. KG, Bremen, Germany) which were located at 5 horizontal locations (−17°, −8.5°, 0°, 8.5°, 17°, vertical midline = 0°; Figure 1). Sound presentation was controlled via a multi-channel soundcard (Creative Sound Blaster Z) and amplified via an audio amplifier (t.amp E4-130, Thomann Germany). Visual stimuli were projected (Acer Predator Z650, Acer Inc., New Taipei City, Taiwan) onto an acoustically transparent screen (Screen International Modigliani, 2×1 m), which was located at 135 cm in front of the participant. The visual stimulus was a cloud of white dots distributed according to a two-dimensional Gaussian distribution (N = 200, SD of vertical and horizontal spread 2°, width of a single dot = 0.12°, duration = 50 ms). Stimulus presentation was controlled using the Psychophysics toolbox (Brainard, 1997) for MATLAB (The MathWorks Inc., Natick, MA) with ensured temporal synchronization of auditory and visual stimuli.

**Figure 1.**
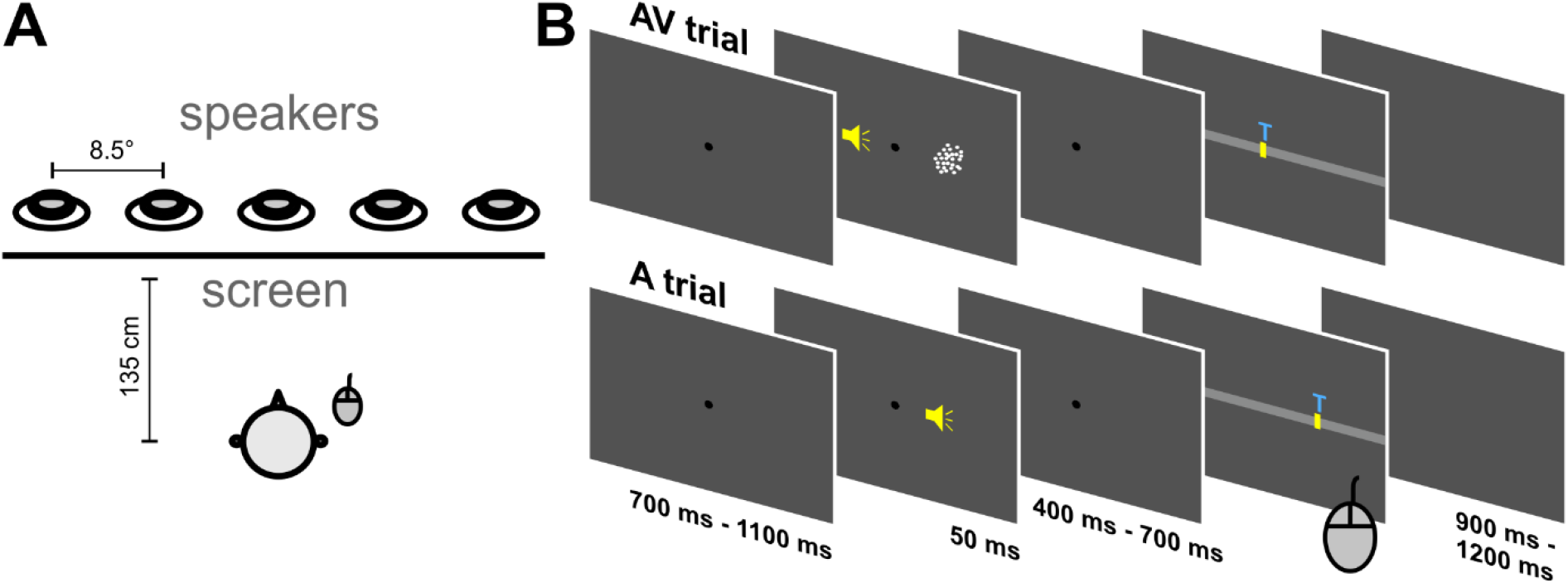
Experimental setup and task. **(A)** Top view of the experimental setup. **(B)** Time-course of the sequence of AV and A trials (V trials are not shown). The yellow speaker icons are for display purposes only. The A trial always followed the AV trial, and participants responded using a mouse.

### 2.3. Experimental Setup

The paradigm was based on a single-trial audio-visual localization task (Park & Kayser, 2019; Wozny & Shams, 2011b), with trials and conditions designed to probe both the ventriloquism effect and the ventriloquism aftereffect. Participants were seated in front of an acoustically transparent screen, with their heads on a chin rest. Five speakers were located immediately behind the screen. Participants responded with a mouse cursor (Figure 1A).

The participants’ task was to localize a sound during either Audio-Visual (AV) or Auditory (A), trials, or to localize a visual stimulus during few Visual (V) trials. We used an auditory report in the multisensory trials in line with previous studies (Bruns & Röder, 2015; Park & Kayser, 2019), and to reduce the length of the experiment in comparison to a dual report paradigm. The locations of auditory and visual stimuli were drawn semi-independently from the 5 locations to yield 9 different audio-visual discrepancies (abbreviated ΔVA in the following; −34°, −25.5°, − 17°, −8.5°, 0°, 8.5°, 17°, 25.5°, 34°). We repeated each discrepancy between the locations of auditory and visual stimuli 44 times for YA, 45 times for OA, resulting in a total of 396 AV-A trial pairs for YA, and 405 for OA. For one OA one block was lost, resulting in 324 pairs for this individual. In addition, on average 72 (YA) and 65 (OA) visual-only trials were interleaved to maintain attention to the visual modality, in line with previous work (Park & Kayser, 2019). Each AV trial was followed by an A trial and the V trials always came after A trials, and hence did not interrupt the AV-A sequence. Trials were pseudo-randomized and divided into 4 blocks for YA and 5 blocks for OA to give these more breaks. Each trial started with a fixation period (uniform 700 ms – 1100 ms), followed by the stimulus (50 ms). After a random post-stimulus period (uniform 400 ms – 700 ms) the response cue emerged, which was a horizontal bar along which participants could move a cursor. A letter ‘T’ was displayed on the cursor for ‘tone’ in the AV or A trials, and ‘V’ for the V trials to indicate which stimulus participants had to localize. There was no constraint on response times. Inter-trial intervals varied randomly (uniform 900 ms – 1200 ms). A typical sequence of trials is depicted in Figure 1B. Participants were asked to maintain fixation during the entire trial except the response, during which they could freely move their eyes.

All participants underwent a test for spatial hearing prior to the main study. We used four of the five potential sound locations (excluding the middle one) and asked participants to indicate the perceived location by pressing left or right keys on a keyboard (2-AFC procedure). The resulting data were fit with a psychometric curve (fitted with a logistic function, free but equal lapse and guess rates) (psignifit toolbox version 4, Schütt et al., 2016) from which we extracted perceptual thresholds (at 50% correct) and slopes.

### 2.4. Model-free analyses of behavioral biases

We defined response biases as follows: The single-trial ventriloquism bias was defined as the bias induced by the visual stimulus away from the true sound location in the AV trial: i.e. the difference between the reported sound location (R_AV_) and the location at which the sound (A_AV_) was actually presented (*ve* = R_AV_ – A_AV_). The single-trial ventriloquism aftereffect was defined as the bias in the reported sound location in the auditory trial relative to the audio-visual discrepancy experienced in the previous trial. It was computed as the difference between the reported sound location (R_A_) and the mean reported location for all A trials of the same stimulus position (μ_RA_), i.e., *vae* = R_A_ – μ_RA_. This was done to ensure that any overall bias in sound localization would not influence this measure, such as a tendency to perceive sounds closer to the midline than they actually are (Rohe & Noppeney, 2015b; Wozny & Shams, 2011b).

We then used generalized linear mixed-effects models (GLMM) to quantify the dependencies of biases on the audio-visual discrepancy (ΔVA), on the response in the AV trial (*R_AV_*), and to test for potential group differences. We used the Bayesian Information Criterion (BIC) to compare the explanatory power of different models, and used the Bayes Factor (BF) based on the BIC to assess the relevance of different model terms (Wagenmakers, 2007).

As a first step, we determined whether each bias is best described as a linear or a non-linear function of ΔVA. A linear dependency is predicted by models of multisensory fusion (Alais & Burr, 2004; Ernst & Bülthoff, 2004), while models of multisensory causal inference additionally predict a non-linear dependency (Cao et al., 2019; Körding et al., 2007; Rohe & Noppeney, 2015b): this non-linear dependency describes the decline in perceptual bias when the audio-visual stimuli are sufficiently discrepant to no longer be judged as originating from a common source. For each bias we determined its dependency on ΔVA by contrasting the predictive power of the following three GLMMs, which were fit across all trial-wise biases of all participants and both groups (using a Gaussian distribution and an identity link function):

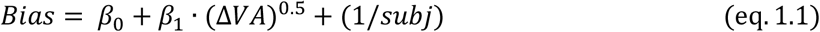

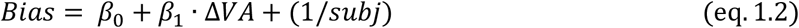

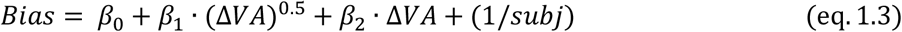

Here *Bias* is either *ve* or *vae,* and *β_1_, β_2_* quantify the magnitudes of the biases and *subj* indexes the participant id as random effects. The square-root term (ΔVA^0.5^) describes the signed square-root of ΔVA (i.e. sign(ΔVA) * sqrt(abs(ΔVA))), and was chosen based on previous work (Cao et al., 2019). Comparing BIC values revealed that model 1.3 provided the best fit for the *ve* and model 1.2 for the *vae*.

For the ventriloquism aftereffect we then asked whether this bias also depends on the previous response (*R_AV_*, localization of the sound in the AV trial). Such a dependency could be expected based on general serial-dependencies seen in many perceptual decision making tasks (Fritsche et al., 2017; Kiyonaga et al., 2017; Talluri et al., 2018), and has been directly suggested to contribute to recalibration biases (Van der Burg et al., 2018). For this we compared model 1.2 to an extended model also including *R_AV_*:

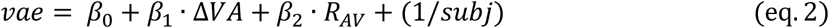

We found that model 2 provided a better fit than model 1.2.

Finally, for both the *ve* and *vae*, we asked whether their dependencies on ΔVA or *R_AV_* differed between groups. We extended models 1.3 (*ve*) and 2 (*vae*) by including the group (G) and its interactions with the discrepancy terms as additional factors:

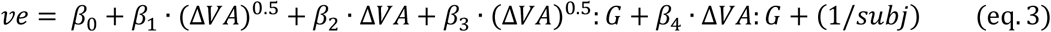

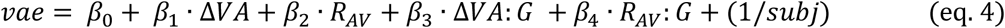

where *G* denotes the group, either YA or OA, and the coefficients *β_3_*, *β_4_* quantify the effect of group on ΔVA or *R_AV_*. To quantify the group-wise influence of these predictors, we also fit models 1.3 (*ve*) and 2 (*vae)* separately for each age group (c.f Table 1, 3).

**Table 1.** Generalized linear mixed-effects models for the ventriloquism effect. **(A)** Model predicting the *ve* based on a linear and non-linear dependency on the multisensory discrepancy (ΔVA), including group as factor (G). **(B)** Group-wise models fit to each group separately. Bayes Factor (BF) were obtained by contrasting the respective full model (models in A or B) against a model omitting the specific factor of interest; a positive BF indicates an improvement in model with by including this predictor; a negative BF indicates that a model without this predictor provides a better fit. CI: 95% confidence interval (parametric).

| (A) $ve \sim \beta_0 + \beta_1 \cdot (\Delta VA)^{0.5} + \beta_2 \cdot \Delta VA + \beta_3 \cdot (\Delta VA)^{0.5} : G + \beta_4 \cdot \Delta VA : G + (1/subj)$ | | | | | | | |
| --- | --- | --- | --- | --- | --- | --- | --- |
| | Name | Estimate ($\beta$) | t-statistics | p-value | CI (95%) | | Log<sub>10</sub>(BF) |
|  | Intercept | -0.4514 | -1.2253 | 0.2205 | -1.1735 | 0.2707 |  |
| | $(\Delta VA)^{\frac{1}{2}}$ | 2.1528 | 6.4496 | 0.0000 | 1.4985 | 2.8071 | 117 |
| | $\Delta VA$ | -0.2577 | -3.7822 | 0.0002 | -0.3912 | -0.1241 | 105 |
| | $(\Delta VA)^{\frac{1}{2}} : G$ | -0.6405 | -3.0183 | 0.0025 | -1.0565 | -0.2246 | 91.8 |
| | $\Delta VA : G$ | 0.2956 | 6.8234 | 0.0000 | 0.2107 | 0.3805 | |
| (B) $ve \sim \beta_0 + \beta_1 \cdot (\Delta VA)^{0.5} + \beta_2 \cdot \Delta VA + (1/subj)$ | | | | | | | |
| | Name | Estimate ($\beta$) | t-statistics | p-value | CI (95%) | | Log<sub>10</sub>(BF) |
| YA | Intercept | -0.7726 | -1.5346 | 0.1249 | -1.7595 | 0.2143 |  |
| | $(\Delta VA)^{\frac{1}{2}}$ | 1.5123 | 10.7897 | < 0.0001 | 1.2375 | 1.7870 | 23.1 |
| | $\Delta VA$ | 0.0379 | 1.3240 | 0.1855 | -0.0182 | 0.0940 | -1.59 |
| OA | Intercept | -0.1155 | -0.2181 | 0.8273 | -1.1532 | 0.9223 |  |
| | $(\Delta VA)^{\frac{1}{2}}$ | 0.8717 | 5.4518 | < 0.0001 | 0.5583 | 1.1852 | 4.48 |
| | $\Delta VA$ | 0.3335 | 10.2164 | < 0.0001 | 0.2695 | 0.3974 | 20.6 |

### 2.5. Statistical Analysis

Confidence intervals (CI, 95%) were obtained using the bootstrap hybrid method with 199 resamples (Bootstrap Matlab Toolbox, Zoubir & Boashash, 1998). Generalized linear mixed-effects models were fit using maximum-likelihood procedures (fitglme.m in MATLAB), using a normal distribution for modelling the response and an identity link function. Bayes Factors were computed as 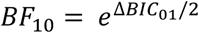, where ΔBIC_01_ = BIC(H_0_) – BIC(H_1_). We compared a model (H_1_) against this model minus a specific predictor of interest (H_0_), therefore BF_10_ > 0 is in favour of the model with the predictor (H_1_), and BF_10_ < 0 vice versa (Wagenmakers, 2007). The magnitudes of BIC differences and BFs were interpreted using established conventions, with BIC differences larger than 6 corresponding to “strong” and those larger than 10 to “very strong” evidence (Kass & Raftery, 1995).

### 2.6. Model-based analysis of the ventriloquism effect

We applied Bayesian causal inference (BCI) models to model the participant-wise ventriloquism biases obtained in the AV trials. These models have been shown to capture the computations underlying flexible multisensory perception and were previously used to investigate ventriloquism-like biases (Cao et al., 2019; Jones et al., 2019; Körding et al., 2007; Rohe et al., 2019; Rohe & Noppeney, 2015b; Wozny et al., 2010). Briefly, the BCI model reflects the inference process about the causal relations of two sensory features based on prior experience and the available sensory evidence, i.e. the presumed observation of the sensory information. Specifically, the model predicts the *a posteriori* probability of a single (C=1) or two distinct (C=2) causes by sampling from a binomial distribution with the common source prior P(C=1) = P_COM_:

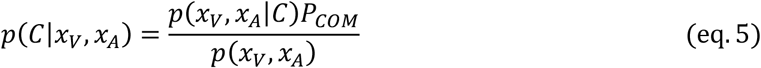

where *x*_*V*_, *x*_*A*_ reflect internal noisy representations of the respective visual and auditory stimuli. For a common source the ‘true’ location (S) is drawn from a Gaussian spatial prior distribution with mean *μ*_*P*_ and standard deviation *σ*_*P*_. For two independent sources the true auditory and visual locations (S_A_, S_V_) are drawn independently from this prior distribution. Sensory noise was modelled by drawing the internal representations from independent Gaussian distributions centred on the true auditory (visual) locations with standard deviations of *σ*_*A*_ and *σ*_*V*_ respectively. Under the assumption of two separate causes, the optimal estimates of the auditory location is given by (Körding et al., 2007):

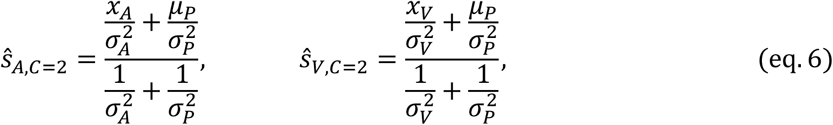

And under the assumption of a common cause the location is determined by the reliability-weighted average (Ernst & Banks, 2002):

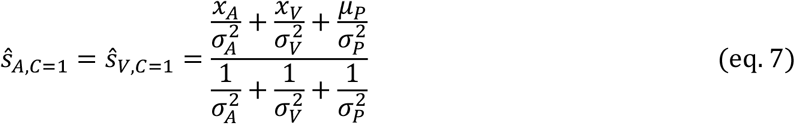

To model a final estimate of the auditory location an observer can combine the estimates under the two causal structures using different decision strategies. We considered three frequently studied strategies (Cao et al., 2019; Körding et al., 2007; Rohe & Noppeney, 2015b; Wozny et al., 2010): Model Averaging (MA), Model Selection (MS), and Probability Matching (PM). With the MA strategy, the final estimate is derived by the weighted-average of the two estimates derived under each causal relation and minimizes the mean expected squared error of the spatial estimates (Körding et al., 2007; Wozny et al., 2010):

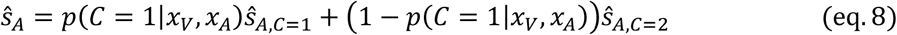

The MS strategy uses the posterior probability to guide the decision by choosing the estimate associated with the causal structure of higher probability:

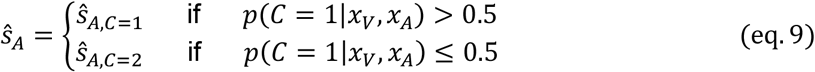

Finally, PM strategy uses a stochastic selection criterion, by choosing a causal scenario in proportion to its posterior probability, which can be implemented by sampling a random ξ from a uniform distribution on each trial:

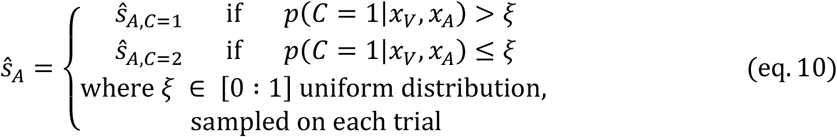

The BCI model for the ventriloquism effect hence comprised the following free parameters, which were determined for each participant and for each decision strategy individually: the uncertainty in the sensory representations *σ*_*A*_, *σ*_*V*_, the width of the prior *σ*_*P*_, the mean of the prior *μ*_*P*_ and the a priori binding tendency P_COM_.

#### 2.6.1. Model Fitting, comparison, and validation

We optimized the model parameters using the BADS toolbox based on the log-likelihood of the true data under the model (Bayesian Adaptive Direct Search (v1.0.3) (Acerbi & Ma, 2017). The predicted conditional distributions for the spatial estimates for each model 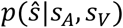 were obtained by marginalizing over the internal variables (*x*_*A*_, *x*_*V*_).

These distributions were obtained by simulating the internal variables 20,000 times for each of the true stimulus conditions and inferring *S*_*A*_ from eq. 8-10. To relate the observed data to the model predictions we binned both into 73 bins (−36° to 36°, increment = 1°), and derived the log-likelihood of each participant’s actual data under a given model from this binned data under the assumption that the different conditions are statistically independent. For each model and participant we repeated the model fitting 500 times and selected the best run with the highest likelihoods. The lower and upper bounds for the initial conditions of the parameters (*σ*_*A*_, *σ*_*V*_, *σ*_*P*_, *μ*_*P*_, *P*_*com*_) were (0.5, 0, 2, −10, 0) and (45, 28, 45, 10, 1), respectively.

We derived the participant specific BIC values from the log-likelihood obtained from the best model fits (BIC = k · ln(N) + 2 · ln(*LL*), k = number of model parameters, N = number of data points to fit model, LL = negative log-likelihood). We derived the group-level BIC value under the assumption that each participant adds independent evidence about the model’s predictive power. To compare the model fit across models and groups, we used the coefficient of determination R^2^ (Nagelkerke, 1991), calculated for each participant and model as 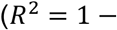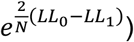. Here LL_1_ denotes the log-likelihood of the fitted model and LL_0_ that of a null model, defined as a model predicting a flat response distribution, and N is the number of data points used to fit the model.

To verify that the model reflects the actual data well, we simulated the predicted behavioral biases from the obtained model parameters for each participant (Palminteri et al., 2017). We simulated 10,000 trials based on the exact stimuli presented to each participant using equations 5-10. We then calculated the mean VE-bias across the simulated trials, deriving a prediction of the average bias under the best participant specific model and parameters (Figure 2).

**Figure 2.**
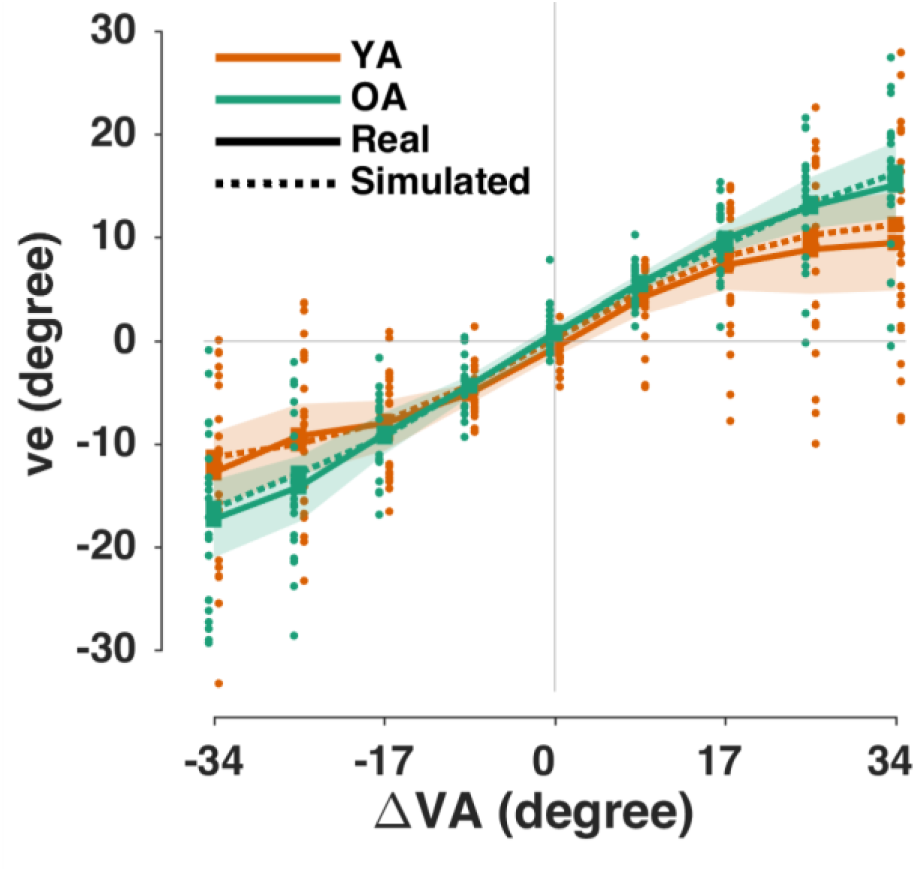
The ventriloquism bias as a function of the audio-visual discrepancy (ΔVA) in the audio-visual trial. Solid lines indicate the mean across participants measured (real) data, shaded areas the 95% hybrid bootstrap confidence interval of the mean, dots indicate single participant data and dotted lines indicate the mean bias predicted (‘simulated’) by the BCI model (see section 2.2). Orange: YA; Emerald: OA.

## 3. RESULTS

### 3.1. Older adults performed worse in a spatial hearing test

All young (YA) and older (OA) participants were devoid of cognitive impairments and had normal vision. Only OA’s with no or mild (< 30dB) hearing loss were included. In addition, all participants performed a spatial hearing test: both groups localized the sounds well and the median accuracy was 97.5% for the YA (range: 85.0% ~ 100%) and 92.5% for the OA (range: 72.5% ~ 100%). Fitting psychometric curves to individual participant’s data revealed no significant group difference in the psychometric threshold reflecting a spatial bias (YA: 1.09° ± 2.56° mean ± SD; OA: 1.00° ± 4.05°; Wilcoxon rank sum test corrected for multiple comparisons with the Holm method p = 0.9112, z value = 0.111). However, the psychometric slope reflecting the sensitivity to spatial information was significantly reduced in the OA (YA: 0.074 ± 0.018, OA: 0.052 ± 0.023, p = 0.01, z value = 2.77).

### 3.2. The ventriloquism bias differs between groups

Both groups exhibited a robust ventriloquism effect for all non-zero discrepancies. Figure 2 shows the ventriloquism bias (*ve*) as a function of the audio-visual discrepancy (ΔVA), and suggests a comparatively smaller bias at larger discrepancies for the YA.

We used GLMMs to test how the ventriloquism bias depends on the multisensory dependency (ΔVA). When tested across both groups this provided ‘very strong’ evidence that the bias is best explained by a model including both a linear and a nonlinear dependency on ΔVA (Table 1, ΔBIC for models 1.1-1.3: 60, 114, 0). This non-linear dependency is in line with predictions from causal inference models, which posit that for large multisensory discrepancies the tendency to bind different stimuli is reduced (Cao et al., 2019; Rohe et al., 2019). To probe for a difference in bias between groups we compared models without and with group as additional factor: this provided ‘very strong’ evidence for an effect of group (ΔBIC without and with group as factor: 414, 0, corresponding to a BF of 91.8). The model coefficients for the model including group as factor revealed significant interactions of group with both dependencies on ΔVA, and the associated BFs provide ‘decisive’ evidence for a group effect (Table 1A). Inspecting the group-wise models (Table 1B) suggests that the ventriloquism bias in the YA is mostly explained by a non-linear dependency on ΔVA, while the bias in the OA depends both in a linear and non-linear manner on ΔV. Importantly, our data show that the non-linear contribution is reduced in the OA compared to the YA, as suggested by the distinct confidence intervals for the respective factor in each group (Table 1B). Hence, the OA fused the audio-visual stimuli over a wider range of discrepancies.

### 3.3. Bayesian inference models reveal no difference in a priori binding tendency

We fit Bayesian causal inference models to individual participant’s data. To assess the quality of the model fit we calculated the coefficient of determination, R^2^ (Nagelkerke, 1991). The models explained the data well (participant-wise best model R^2^ mean ± SEM: YA = 0.88 ± 0.017, 95%-CI = [0.84, 0.91]; OA = 0.76 ± 0.045, CI = [0.67, 0.88]). Using model validation we confirmed that the model was able to reproduce the average bias for individual participants (Figure 2, dotted lines). Yet, a non-parametric permutation test (two-sided, exchanging participant-wise group-labels) showed that the group-averaged model fit differed significantly between groups (N = 50000, p = 0.009).

Inspecting the participant-wise best fitting models revealed that the decision strategy best explaining the observed data was heterogeneous among participants in both groups (Fig. 3A). However, the group-wise BIC values indicated that for the YA a model based on probability matching (PM), and for the OA a model based on model selection (MS), provided the overall best fit (relative summed-BIC values YA: [640, 419, 0] for [MA, MS, PM]; OA: [491, 0, 38]). Hence despite the overall heterogeneity our data suggest that the preferred decision strategies could possibly differ with age.

**Figure 3.**
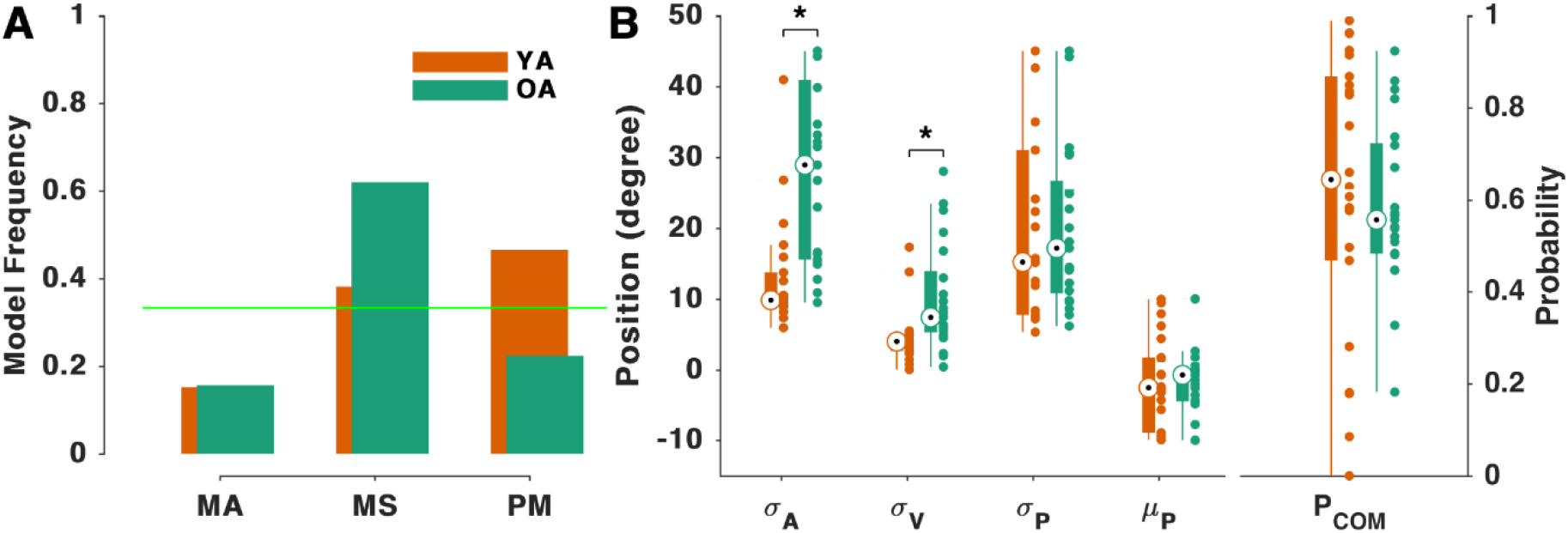
Modelling the ventriloquism effect using Bayesian causal inference. **(A)** Model frequencies for the three potential decision strategies obtained from participant-specific best fitting models. MA: model averaging, MS: model selection, PM: probability matching. **(B)** Model parameters for individual participants’ best model. Black circles denote the median, and the bars are the 25% and 75% percentiles. Dots are individual participants. Asterisks denote a difference in the model parameters between groups (p < 0.01, Rank sum test). Orange: YA; Emerald: OA.

The distribution of the obtained participant-specific best-fitting model parameters (Fig. 3B) matches previous reports obtained using similar models and paradigms (Jones et al., 2019; Rohe & Noppeney, 2015b; Wozny & Shams, 2011a). Comparing the model parameters between groups revealed a significantly reduced sensory precision for auditory and visual spatial information in the OA (Figure 3B; see Table 2 for statistical tests), matching the reduced sensitivity for spatial sounds in the pre-test (see above). The other model parameters did not differ significantly, including the *a priori* belief that the auditory and visual stimuli are causally linked (P_COM_) (Table 2). In particular, the confidence intervals for P_COM_ largely overlapped between groups. The model-based analysis hence reveals differences in the precision of unisensory estimates, but also shows that both groups tended to combine the multisensory evidence based on a comparable a priori belief in a common source.

**Table 2.**
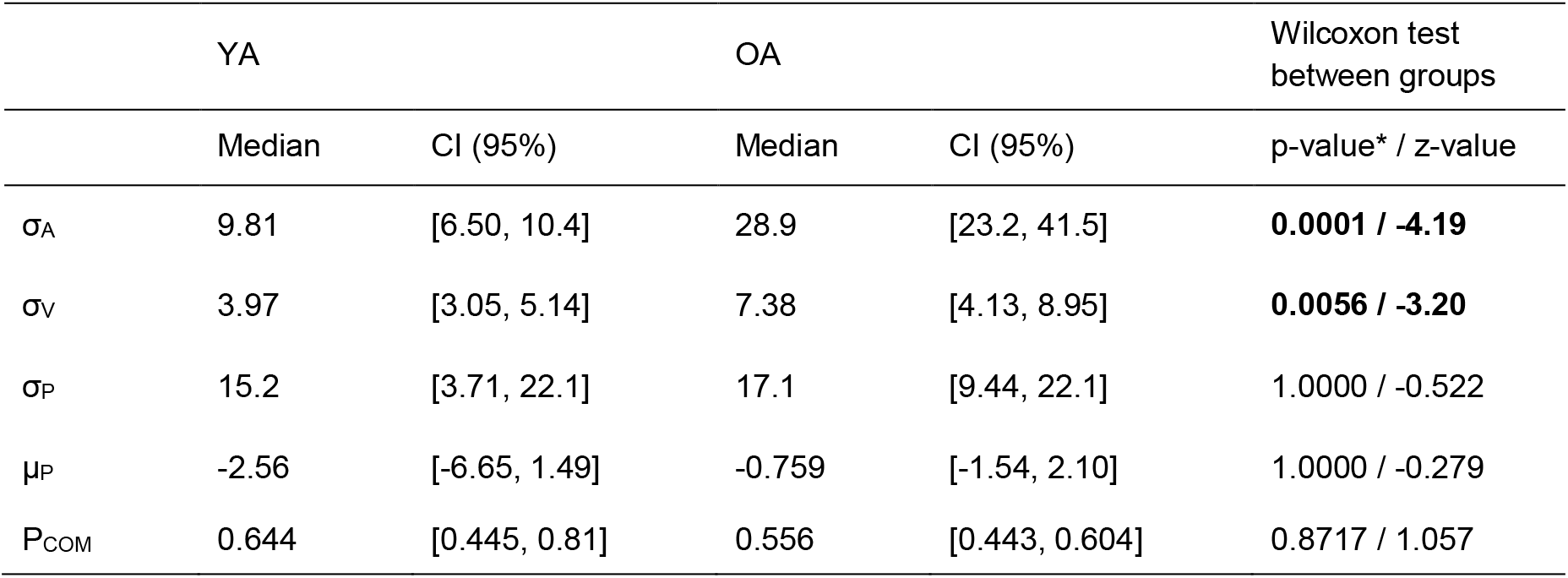
Model parameters of participant-specific best-fitting Bayesian causal inference models. YA: younger adults, OA: older adults. σ_A_: standard deviation of the auditory likelihood, σ_V_: standard deviation of visual likelihood, σ_P_: standard deviation of prior, μ_P_: mean of prior, P_COM_: a priori binding tendency. *corrected for multiple comparisons with Holm’s method. CI: 95% confidence interval based on bootstrapping.

### 3.4. The drivers of the ventriloquism aftereffect differ between groups

Both groups exhibited a robust ventriloquism aftereffect bias (*vae*), shown in Figure 4 as a function of the audio-visual discrepancy.

**Figure 4.**
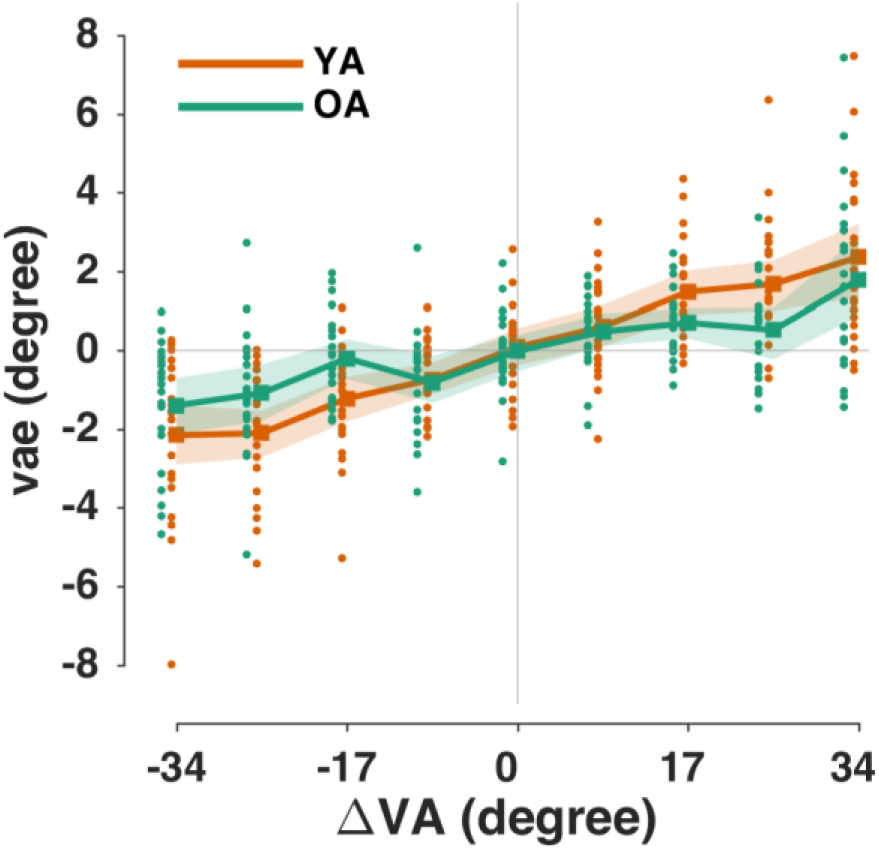
The ventriloquism aftereffect bias (obtained in the auditory trial) as a function of the audio-visual discrepancy (ΔVA; presented in the audio-visual trial). Solid lines indicate the mean across participants, dots indicate single participant data and shaded areas the 95% hybrid bootstrap confidence interval of the mean. Orange: YA; Emerald: OA.

Following the same strategy as for the ventriloquism effect, we first determined the predictors necessary to model the single-trial ventriloquism aftereffect across all participants and both groups. Comparing the three candidate GLMMs in eq. (1) provided ‘strong’ evidence that the aftereffect bias is best explained by a model including only a linear dependency on discrepancy (ΔBIC for models 1.1-1.3: 10, 0, 9). In addition, we found ‘very strong’ evidence that adding the previous response (R_AV_) as additional factor further improved the model fit (ΔBIC without 174 and with R_AV_ as factor 0; BF = 16), as expected based on work on serial dependencies in perceptual decision in general (Fritsche et al., 2017; Kiyonaga et al., 2017; Talluri et al., 2018). Importantly, we then tested whether adding the group as additional factor (eqs. 2 and 4) further improved the model fit and found ‘very strong’ evidence for an improved fit (ΔBIC without 414 and with group as factor 0; BF in favour of the group model 10.7). The resulting coefficients of the full model (eq. 4) revealed a significant interaction of the factors group with discrepancy (ΔVA) and with the previous response (R_AV_) (Table 3A), and the associated BFs provide ‘decisive’ evidence for group effects.

**Table 3.** Generalized linear mixed-effects models for the ventriloquism aftereffect. (**A)** Model predicting the *vae* based on the multisensory discrepancy (ΔVA), the previous response (R_AV_) and their interactions with the group factor (G). **(B)** Group-wise models fit to each group separately. Bayes Factor (BF) were obtained by contrasting the respective full model (models A or B) against a model omitting the specific factor of interest; a positive BF indicates an improvement in model with by including this predictor; a negative BF indicates that a model without this predictor provides a better fit. CI: 95% confidence interval (parametric).

| **(A) $vae \sim \beta_0 + \beta_1 \cdot \Delta VA + \beta_2 \cdot R_{AV} + \beta_3 \cdot \Delta VA : G + \beta_4 \cdot R_{AV} : G + (1/subj)$** |  |  |  |  |  |  |
| --- | --- | --- | --- | --- | --- | --- |
| | Name | Estimate ($\beta$) | t-statistics | p-value | CI (95%) | Log<sub>10</sub>(BF) |
|  | Intercept | 0.0450 | 0.7958 | 0.4262 | -0.0659 0.1560 |  |
| | $\Delta VA$ | 0.1234 | 13.9894 | < 0.0001 | 0.1061 0.1407 | 128 |
| | $R_{AV}$ | 0.0005 | 0.0341 | 0.9728 | -0.0275 0.0285 | 26.7 |
| | $\Delta VA : G$ | -0.0396 | -7.1401 | < 0.0001 | -0.0505 -0.0288 | 10.7 |
| | $R_{AV} : G$ | 0.0302 | 3.3490 | 0.0008 | 0.0125 0.0479 | |
| **(B) $vae \sim \beta_0 + \beta_1 \cdot \Delta VA + \beta_2 \cdot R_{AV} + (1/subj)$** |  |  |  |  |  |  |
| | Name | Estimate ($\beta$) | t-statistics | p-value | CI (95%) | Log<sub>10</sub>(BF) |
| YA | Intercept | 0.0436 | 0.6152 | 0.5384 | -0.0952 0.1823 |  |
| | $\Delta VA$ | 0.0838 | 23.7736 | < 0.0001 | 0.0769 0.0907 | 117 |
| | $R_{AV}$ | 0.0307 | 5.3749 | < 0.0001 | 0.0195 0.0419 | 4.29 |
| OA | Intercept | 0.0466 | 0.5244 | 0.6000 | -0.1275 0.2206 |  |
| | $\Delta VA$ | 0.0441 | 10.2827 | < 0.0001 | 0.0357 0.0525 | 20.9 |
| | $R_{AV}$ | 0.0609 | 8.6451 | < 0.0001 | 0.0471 0.0748 | 14.2 |

Inspecting the group-wise models (Table 3B) revealed that the slopes for the factors ΔVA and R_AV_ had comparable magnitude in the OA, while for the YA the dependency on the sensory discrepancy ΔVA was more than twice as strong as the dependency on R_AV_. Furthermore, the confidence intervals for each predictor and group suggest that the aftereffect in the OA is more dependent on the previous response, while the aftereffect in the YA is more dependent on the multisensory discrepancy. Hence, the present data suggest a differential dependency of the aftereffect on previous sensory and previous motor-related signals in two groups.

The ventriloquism aftereffect bias is a direct perceptual consequence of the ventriloquism effect (Lewald, 2002; Radeau & Bertelson, 1974). Therefore, we tested for a participant-specific correlation between the single-trial *ve* and *vae* biases. This trial-wise coupling was significant for 18 out of 22 participants in the YA, and 16 out of 21 in the OA (Spearman correlation at p < 0.05). The median correlation coefficients were 0.201 and 0.202 for YA and OA respectively, and a permutation test suggested that the values did not differ between the two groups (50000 permutations of group-labels, p = 0.9670). Hence, the effective trial-by-trial correlation of the two multisensory responses biases was comparable between young and old participants.

## 4. DISCUSSION

We used an audio-visual spatial localization task to characterize age-related changes in the within-trial and between-trial ventriloquism effects. In our data both the ventriloquism bias and the immediate aftereffect differed between groups. Our results suggest that the age-difference in the ventriloquism bias can be largely attributed to changes in peripheral spatial hearing rather than a change in sensory causal inference, while the age-difference in the aftereffect can be attributed to a shift from a sensory-driven to a behavior-driven influence of past experience on subsequent choices with increasing age.

### 4.1. Changes in within-trial multisensory integration with age

Previous studies reported a number of changes in multisensory perception with age. For example, OA seemed to benefit more from multisensory compared to unisensory information during speeded detections (Laurienti et al., 2006; Mahoney et al., 2014; Zou et al., 2017), had longer temporal binding windows (Basharat et al., 2018; Chan et al., 2014b; DeLoss et al., 2013; Hay-McCutcheon et al., 2009; Noel et al., 2016), experienced increased difficulty to segregate multisensory stimuli (Setti et al., 2011), and more strongly fused audio-visual speech (Sekiyama et al., 2014; Setti et al., 2013). Contrasting this view, recent work suggests that healthy young and aging brains may follow similar computational rules when combining multisensory information (Billino & Drewing, 2018; Campos et al., 2018; Cressman et al., 2010; Jones et al., 2019), except possibly that the OA may take longer to respond as a result of age-related slowness (Jones et al., 2019. see also Smith & Brewer, 1995; Starns & Ratcliff, 2010).

In line with studies reporting age-related changes in multisensory perception, we found a stronger non-linear influence of multisensory discrepancy on the ventriloquism bias in the YA. In contrast, the bias in the OA revealed a more linear dependency. The combined evidence from the spatial hearing test and Bayesian modelling suggests that the stronger and more linear bias in the OA effectively results from a loss in spatial hearing rather than a change in the *a priori* belief in a common source for the audio-visual stimuli. Our results hence suggest the loss of precision in peripheral sensory representations as a major driver of age-changes in multisensory behaviour. Such a loss in sensory precision reduces the influence of the affected modality on multisensory behaviour, resulting in a stronger and more widespread (over discrepancies) bias induced by another modality on perceptual judgements in the affected modality. Such a peripheral cause can also explain why a recent study reported no difference in spatial ventriloquism biases between younger and older participants (Jones et al., 2019). In that study the older participants had comparable spatial hearing to the YA (Jones et al., 2019).

Bayesian causal inference models allow separating two potential origins for the age-change in ventriloquism bias: a peripheral loss in sensory acuity and a change in how cognitive processes combine disparate sensory information for behaviour (Körding et al., 2007; Rohe & Noppeney, 2015a; Wozny et al., 2010). We found that models fit to OA’s responses featured larger uncertainties in their sensory representations but exhibited comparable a priori believes into a common source for the multisensory stimuli. Hence, our results clearly speak towards a reduced peripheral acuity in the OA and confirm a previous study, which found no difference in the a priori binding tendency with age in healthy individuals (Jones et al., 2019). Hence, despite changes in neurophysiological markers of sensory encoding and cognition as well as changes in general brain structure with age (Henry et al., 2017; Henschke et al., 2018; McNair et al., 2019) the converging evidence suggests that multisensory perception in healthy aging is mostly affected by changes in peripheral sensory processes.

The typical BCI models considered in the literature comprise a number of decision strategies (Körding et al., 2007; Rohe & Noppeney, 2015b; Wozny et al., 2010). While the individual-participant data were heterogeneous, the group-level quality of fit suggested a difference between groups: for the YA probability matching provided the best fit, in line with previous studies on spatial perception in young participants (Wozny et al., 2010; Wozny & Shams, 2011a). In contrast, for the OA model selection was the best strategy. Hence, in addition to peripheral sensory processes the decision strategies used to convert sensory evidence into an overt response may also change with age. Yet, further work is required to confirm this hypothesis and investigate it in the context of more general changes in cognitive strategies with age (Dully et al., 2018; Koen & Rugg, 2019; Nyberg et al., 2012; Roberts & Allen, 2016).

### 4.2. Changes in between-trial multisensory recalibration with age

Similarly as for within-trial multisensory integration, age-changes have also been reported for multisensory recalibration paradigms (Buch, 2003; Chan et al., 2014a): for example, OA were found to show reduced visuo-motor adaptation (Buch, 2003) and reduced adaptation to audio-visual synchrony (Chan et al., 2014a). Yet, often the results were mixed (Buch, 2003). Our data reveal a robust trial-by-trial ventriloquism aftereffect in both groups, which was comparably linked (correlated) to the respective preceding single trial ventriloquism bias in both groups. However, the driving factor for this aftereffect seems to change with age: while for the YA the aftereffect was more influenced by the previous multisensory discrepancy, for the OA this bias featured a prominent contribution from their previous motor response.

Serial dependencies in perceptual decisions can be driven by multiple factors, such as the previous stimuli (Bruns & Röder, 2015; Kayser & Kayser, 2018; Park & Kayser, 2019; Wozny & Shams, 2011b), the previous responses (Fritsche et al., 2017; Kiyonaga et al., 2017; Talluri et al., 2018), or a meta-cognitive assessment of previous behaviour (Benwell et al., 2019). Yet, previous work in young participants considered the ventriloquism aftereffect generally as a sensory-driven phenomenon (Park & Kayser, 2019; Van der Burg et al., 2018). Confirming this view, we found that the sensory contribution to the aftereffect prevailed in the YA. However, the previous response carried significant predictive power for the aftereffect in both groups, and for the OA, the previous response contributed substantially more to the bias than the multisensory discrepancy.

How can we understand this shift from a sensory-driven to a behavior-driven influence of past experience on multisensory behaviour with increasing age? Previous work has linked the trial-by-trial ventriloquism aftereffect to brain structures implied in memory (Park & Kayser, 2019). While general memory declines with age (Nyberg & Pudas, 2019), episodic memory seems to be more affected than procedural memory (Nyberg et al., 2012; Small, 2001). The active nature of the overt response in the present paradigm (cursor movement) may generally boost its maintenance over time compared to the sensory information (Cohen, 1989), introducing a prominent procedural component in the aftereffect bias. On top of this, the reduced sensitivity to auditory spatial information may have further shifted the dependency from previous sensory to motor information in the OA. Hence, the shift from a sensory-based adaptive bias to a motor-based bias with age may be a multi-faceted effect arising from both a reduced episodic memory and a reduced sensory precision with age.

### 4.3. Conclusion

We found that two frequently studied multisensory response biases change with age: the audio-visual ventriloquism effect and its trial-by-trial aftereffect on subsequent auditory localization. Our collective evidence suggests that the age-effect in the ventriloquism bias results from a loss in peripheral precision rather than a change in causal inference processes typically associated with parieto-frontal brain function. The age-difference in the aftereffect may in part also be driven by a loss in peripheral function, but our data also reveal the emergence of a prominent motor-related component driving this response bias with age, pointing to compensatory memory-related processes that influence multisensory behaviour in the elderly. Given the emerging understanding of the brain structures underlying multisensory causal inference and recalibration (Cao et al., 2019; Körding et al., 2007; Park & Kayser, 2019; Rohe & Noppeney, 2015b), future neuroimaging studies could now confirm whether the sensory computations underlying multisensory behaviour and their neural implementations are largely preserved with age, even when peripheral sensory processes decline.

## ACKNOWLEDGEMENTS

This work was supported by the European Research Council (to C.K. ERC-2014-CoG; grant No 646657).

